# Tau fibrils induce nanoscale membrane damage and nucleate cytosolic tau at lysosomes

**DOI:** 10.1101/2023.08.28.555157

**Authors:** Kevin Rose, Tyler Jepson, Sankalp Shukla, Alex Maya-Romero, Martin Kampmann, Ke Xu, James H. Hurley

## Abstract

The prion-like spread of protein aggregates is a leading hypothesis for the propagation of neurofibrillary lesions in the brain, including the spread of tau inclusions associated with Alzheimer’s disease. The mechanisms of cellular uptake of tau seeds and subsequent nucleated polymerization of cytosolic tau are major questions in the field, and the potential for coupling between the entry and nucleation mechanisms has been little explored. We found that in primary astrocytes, endocytosis of tau seeds leads to their accumulation in lysosomes. This in turn leads to lysosomal swelling, deacidification and recruitment of ESCRT proteins, but not Galectin-3, to the lysosomal membrane. These observations are consistent with nanoscale damage of the lysosomal membrane. Using live cell and STORM, imaging, nucleation of cytosolic tau occurs primarily at the lysosome membrane under these conditions. These data suggest that tau seeds escape from lysosomes via nanoscale damage rather than wholesale rupture, and that nucleation of cytosolic tau commences as soon as tau fibril ends emerge from the lysosomal membrane.

---

Neurodegenerative diseases of protein misfolding are characterized by the abnormal aggregation of host proteins into amyloid inclusions. These inclusions are defined by the presence of a cross-β structural core ^1^. The major neurodegenerative diseases, including Alzheimer’s disease (AD), Parkinson’s disease (PD), and amyotrophic lateral sclerosis (ALS), are characterized by progressive spread of these protein aggregates within the brain ^2^. The leading model for spread is that inclusions propagate by a prion-like mechanism, in which seeds spread from donor cells and nucleate the misfolding and aggregation of endogenous protein in recipient cells ^3,4^.

One such class of prion-like neurodegenerative diseases, termed “tauopathies”, is associated with the aggregation of the protein tau. Tau is a microtubule-associated protein encoded by a single gene but alternatively spliced into six expressed isoforms in humans, which vary in the number of N-terminal domains (N) and microtubule-binding repeats (R) ^5^. Tau aggregation is considered to be a driver of AD and also a larger family of neurodegenerative diseases known as tauopathies ^5^. Tauopathies are diagnosed by identification of tau protein aggregates in neurons, astrocytes and oligodendrocytes of post-mortem brain tissue ^6–9^. In general, the structure of tau fibrils is polymorphic, and each tauopathy is characterized by a unique fibril structure as revealed by high-resolution cryo-EM ^10,11^.

Tau is thought to spread between cells when seeds are taken up in recipient cells via endocytosis ^12^. Once seeds have entered recipient cells, tau seeds are capable of nucleating cytosolic tau aggregation ^13–15^. The pathway and egress of tau seeds from the endocytic and endolysosomal system has been a major question in the field. One leading hypothesis has been that tau seeds escape via lysosomal rupture. Galectins have been used as a marker for rupture because they bind to the newly exposed lumenal glycans of ruptured lysosomes and trigger their degradation via lysophagy ^16^. Previous studies conducted in immortalized and primary cells have demonstrated lysosomal accumulation of tau fibrils with evidence of lysosome damage occurring using a Galectin-based reporter system in some reports ^17,18,19^, although not in others ^14,15^. The inconsistency in these reports suggests that lysosomal rupture, at least on the scale necessary to expose lumenal glycans, cannot be a general explanation for tau escape.

It was recently reported that ESCRT proteins, specifically the ESCRT-III subunit CHMP6, suppressed tau fibril escape from the lysosomal lumen to the cytosol ^14^. The ESCRT proteins are a conserved machinery for membrane budding and sealing ^20,21^ in many pathways in cells, including ectosome and virus release, cytokinetic abscission, multivesicular body biogenesis in the endolysosomal pathway, and membrane repair. Among their many functions, ESCRTs are responsible for repairing nanoscale lysosomal membrane damage ^22–25^. Damage sensing is thought to be mediated by the leakage of Ca^2+^ from the lysosome as detected by ALG-2 and other Ca^2+^-and membrane-binding proteins ^22,23,26^. This form of membrane damage is distinct from the larger ruptures detected by Galectin-3 and triggers a repair response rather than lysophagy ^24,25^. Consistent with a potential role for nanoscale lysosomal membrane damage in tau escape, the lysosome membrane damaging agent leucyl-leucyl-O-methyl ester (LLOME) accelerates seeded tau aggregation ^14^. Nevertheless, ESCRTs act at multiple points in both the biogenesis and repair of lysosomes, and the crucial question of the mechanism of lysosomal escape by tau seeds remained to be determined.

Here, we set out to determine if tau does in fact escape from lysosomes via nanoscale membrane damage. We found that tau fibril accumulation within human primary astrocyte lysosomes triggered membrane permeabilization and an increase in lysosomal pH that was sensed by ESCRTs. Using confocal and STORM microscopy, we characterized the ultrastructure of the fibril-burdened lysosomal compartment and identified lysosomes with tau fibrils protruding from their membranes. These findings reveal a direct mechanism where tau seeds induce lysosomal membrane permeabilization and trigger the templated aggregation of cytosolic tau at the surface of the lysosome.

## Results

### Tau fibrils deacidify lysosomes

We established a primary human astrocyte-based model of endocytosis of pre-formed 4R tau fibrils (PFF), which we used to monitor the effect of fibril uptake on the endolysosomal system. In astrocytes, tau accumulation induces the formation of astrocytic plaques and tufted astrocytes, the diagnostic criterion for Corticobasal Degeneration (CBD) and Progressive Supranuclear Palsy (PSP), respectively ^8,9^. Tau inclusions in astrocytes are typically composed of a truncated form of tau containing 4 C-terminal repeats (4R tau) ^27^, which was therefore used to generate fibrils in this study. Astrocytes acquire extracellular tau by a variety of mechanisms ^7,18,28–31^. We therefore carried out the study in astrocytes as a system directly relevant to disease, robust in its response, and tractable for the imaging experiments described below.

The hypothesis that tau fibrils induced nanoscale damage to the lysosomes predicts that exposure to fibrils should create leaks that increase lysosomal pH. To test for changes in pH, normal human brain-derived astrocytes were transduced with a lysosome activity reporter (pHLARE) which monitors lysosomal pH (Figure 1a). pHLARE is a late endosome associated protein-1 (LAMP1)-derived construct that is a single-pass transmembrane protein with tandem fusions to lumenal pH-sensitive GFP and cytosolic RFP ^32^. Healthy lysosomes exhibit a pH of ∼5, but loss of a proton gradient due to membrane permeabilization or impairment of the lysosomal proton pumping V-ATPase can drastically raise the pH to ∼7, equal to that of the cytosol ^32^. At rest, approximately 14% ± 15% of astrocyte lysosomes exhibited detectable GFP levels. Treatment with 250 nM Bafilomycin A1 (BafA1) for 6 hours, an inhibitor of the V-ATPase, increased the number of lysosomes with high pH to 47% ± 17%. Similarly, although to a lesser extent, incubation with 250 nM PFF for 48 hours increased the number of high pH lysosomes to 35% ± 18% (Figure 1b).

**Figure 1:**
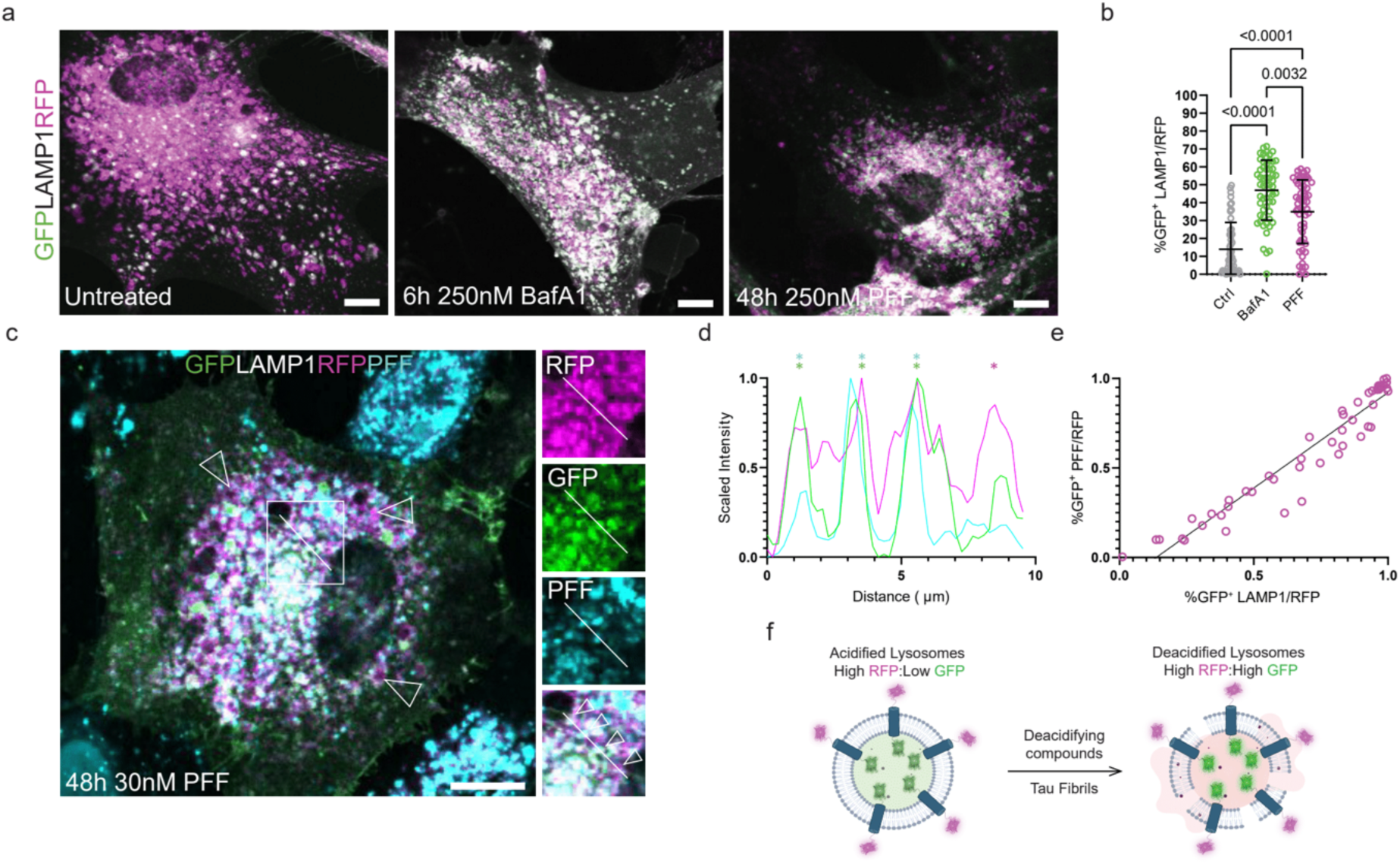
Tau fibrils permeabilize lysosomes. Primary human astrocytes were stably transduced with a GFP-LAMP1-RFP tandem fusion protein to stoichiometrically monitor lysosomal pH. At rest, most lysosomes were RFP-positive. After treatment with Bafilomycin or PFF, the fraction of GFP-positive lysosomes increased dramatically (a). Quantification and 2way ANOVA of total cell GFP area normalized to total RFP signal prior to analysis (b) (n=52, 58, and 60 cells for untreated, BafA1, and PFF conditions, respectively, individual cells plotted with mean and standard deviation). The fibril seeding assay was performed in parallel using 30 nM fluorescently labeled PFF (c). A representative line scan from a cell with four neighboring lysosomes and only one containing labeled fibrils (RFP in magenta, GFP in green, and fibrils in cyan) (d). (e) Line plot revealing a positive linear correlation between cells that had GFP-positive lysosomes and GFP-positive PFF (R^2^=0.9215, n=50 cells, plotting individual Manders co-localization co-efficients). Schematic detailing the effect of deacidifying compounds and tau fibrils on the state of lysosomal pH (f). Scale bar: 20 µm.

To determine if the observed increase in lysosomal pH induced by PFF endocytosis correlated with the presence of PFF in the lysosomal lumen, we repeated our seeding assay with fluorescently labeled PFF. In these experiments, the concentration of PFF added to the media was 30 nM, which is within 3-fold of reported CSF levels of tau in patient brains (Figure 1c) ^33^. At this near-physiological concentration of tau seeds, we could identify neighboring lysosomes in the same cells that demonstrated clear fibril-dependent pH differences (Figure 1d). We also detected a clear positive correlation between the lumenal presence of PFF and deacidified lysosomes (Figure 1e). These data show that burdening the endolysosomal system with PFF potently neutralizes lysosomal pH (Figure 1f).

### ESCRTs, but not Galectins, are recruited to PFF-containing lysosomes

Having observed a clear effect of PFF on the ionic state of the lysosomal compartment, we suspected that PFF could be either interfering with V-ATPase function ^34,35^, or mechanically damaging lysosomal membranes leading to proton leakage. To test for a lysosome damage response to PFF, we first transduced human primary astrocytes with TMEM192-RFP to label lysosomes, and treated cells with fluorescently labeled PFF. PFF accumulated in the lysosomal compartment labeled by TMEM192-RFP, in agreement with previous reports ^14,15^ (Figure 2a). We next transduced TMEM192-RFP expressing astrocytes with CHMP4B, a core component of the ESCRT system ^20,21^, fused to a c-terminal HALO tag with a long linker known to support function ^36^ to test for ESCRT recruitment after PFF treatment. At rest, CHMP4B expression was diffuse with few puncta and did not interact with TMEM192-positive lysosomes (Figure 2b). After fibril treatment, CHM4B formed large puncta on the surface of lysosomes and PFF, consistent with a triggered nanoscale damage repair response (Figure 2c-e). Consistent with other recent reports ^14,15^, we did not observe a significant increase in Galectin-3 recruitment to endogenously labeled lysosomes, suggesting that PFF permeabilize, but do not rupture the lysosomes of astrocytes (Figure 2f-h). Substantial numbers of Galectin-3 puncta are present in both control and PFF-treated cells, showing that a significant level of lysosomal rupture already exists in the absence of treatment. These data show that in astrocytes, tau fibrils permeabilize but do not rupture lysosomes.

**Figure 2:**
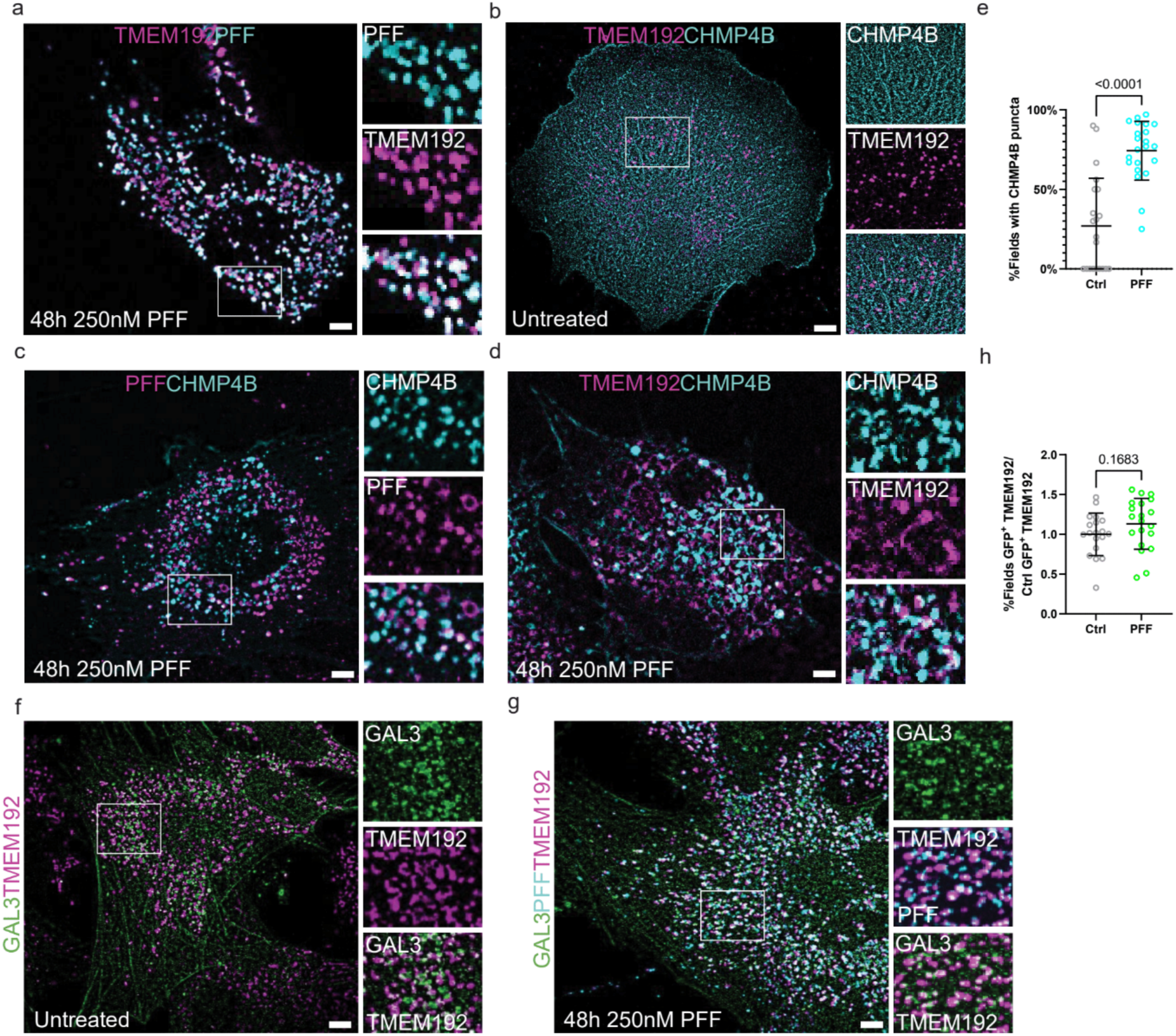
CHMP4B, but not Galectin-3 is recruited to fibril-containing lysosomes. TMEM192-RFP expressing astrocytes were treated with labeled PFF to verify endosomal uptake (a). These same cells were additionally transduced with CHMP4B-HALO to monitor potential ESCRT recruitment to fibril-containing lysosomes. At rest, CHMP4B was mostly diffuse with few puncta that rarely interacted with TMEM192-positive lysosomes (b). Treatment with labeled PFF triggered the robust formation of CHMP4B puncta that overlapped with fibrils and lysosomes (c-d). (e) Quantification of CHMP4B-expressing cells before and after PFF treatment (n=3 replicates, 20 random and independent fields of 10-20 cells per replicate, n=386 cells, represented as mean and standard deviation across fields). TMEM192-RFP expressing astrocytes were transduced with Gal3-GFP and the fractional overlap of GFP puncta area and endogenous TMEM192-RFP area was compared before and after fibril treatment (f-h) (n=4, 20 random and independent fields of 10-20 cells per replicate, n=372 cells). Scale bars: 10 µm.

### Soluble tau aggregates appear adjacent to lysosomes

Although tau fibril endocytosis has been well documented, it is still unclear whether lysosome-associated fibrils are the species that trigger cytosolic tau aggregation ^14,15^ (Figure 3a). To address this question, we applied the astrocyte model of PFF endocytosis to study the effect of fibril uptake on the state of cytosolic tau (Figure 3b). First, we transduced our TMEM192-RFP-expressing astrocytes with a mClover2-tagged truncated version of aggregation-sensitive tau (K18.tau) ^12,14^. In the absence of fibrils, K18.tau was diffuse and/or localized to stress fibers, and did not specifically interact with lysosomes or microtubules (Figure 3b).

**Figure 3:**
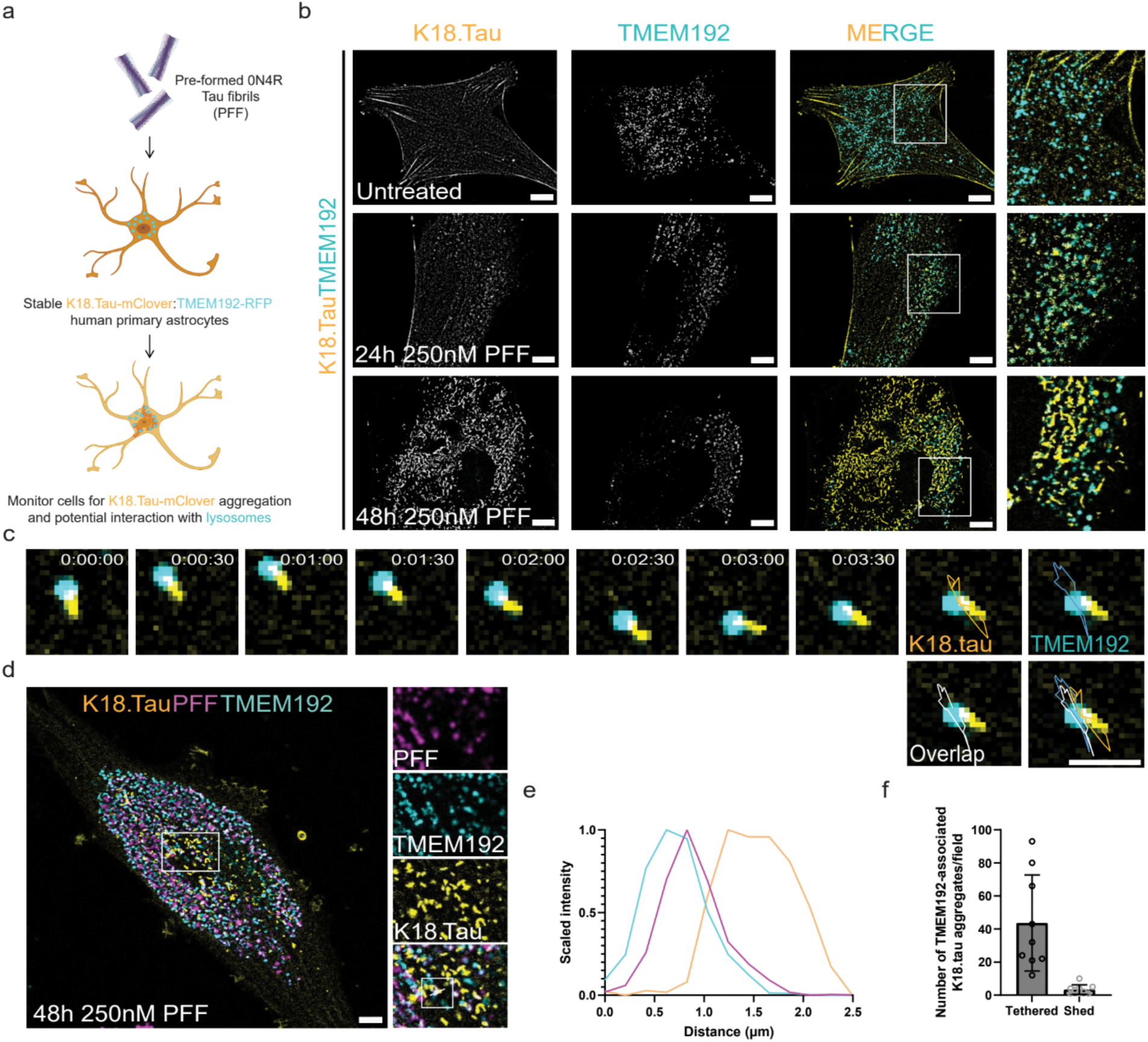
Soluble tau aggregation occurs in the proximity of lysosomes. To model the tau fibril uptake mechanism by astrocytes in vitro, we treated astrocytes stably expressing K18.tau-mClover and TMEM192-RFP with PFF (a). These dual astrocytes were used in live imaging experiments to determine what impact PFF endocytosis had on the cytoplasmic pool of tau (b). At rest, cells had diffuse K18.tau expression that did not interact with lysosomes (top panel). After 24 hours of fibril treatment, K18.tau aggregates could be detected in close proximity to TMEM192-positive lysosomes (middle panel). The amount of K18 aggregation increased at 48 hours post-treatment with numerous lysosomes found associated with large K18 aggregates (lower panel). Representative trace of a lysosome-associated K18 aggregate during brief time-lapse imaging (c) (n=393). Representative cell from the PFF seeding assay repeated with fluorescently labeled PFF to verify lysosomal accumulation and association with K18 aggregates (d) (n=100). Fluorescence intensity plot of the line scan shown in (d), revealing the spatial distribution of PFF, lysosomes, and K18 aggregates (e). (f) Quantification of the number of lysosome-tethered versus shed tau aggregates (n=3 replicates, 3 random and independent fields of 20 cells per replicate, n=60 cells). Scale bars: 10 µm (b,d); 2 µm (c).

The dual reporter astrocytes were then treated with 250 nM of PFF and monitored for aggregation for 1-2 days. K18.tau aggregates appeared adjacent to lysosomes within 24 hours of incubation with fibrils (Figure 3b). At 48 hours post fibril uptake, K18.tau aggregates appeared to be elongated, resembling the tau tangles from those of tufted astrocytes ^37^ and interacted dynamically with lysosomes (Figure 3c). Approximately 93% of lysosome-associated elongated K18.tau aggregates remained tethered to the lysosome surface over a three-minute window of observation (Figure 3c-d). Imaging labeled PFF with K18.tau showed that the K18.tau aggregates extended from the exterior of lysosomes containing PFF (Figure 3e-f). These data strongly suggest that lysosomal PFF are responsible for nucleating the observed cytosolic tau aggregates. The data described thus far suggest that tau fibrils can permeabilize lysosomes, thereby exposing fibril ends to the cytosol and triggering aggregation of cytosolic tau in the vicinity of lysosomes (Figure 3g).

### Spatial organization of tau aggregates on the surface of fibril-burdened lysosomes

In order to resolve more precisely the spatial relationship between lysosomal PFF and cytosolic K18.tau aggregation, we employed multicolor STORM super-resolution microscopy ^38^. Primary human astrocytes were transduced with the K18.tau-mClover fibril-induced aggregation sensor, and treated with 647-labeled PFF for 48 hours. Cells were stained for endogenous LAMP1, which was detected using a fluorescent secondary antibody with a CF680 label, and K18.tau-mClover was stained using anti-GFP antibodies with a CF583R secondary antibody. In the absence of PFF, there was no detectable GFP aggregagtion on the surface of LAMP-1-positive lysosomes. However, upon introduction of PFF into the media, fibril accumulation within LAMP-1-positive vesicles was readily apparent, and GFP aggregates were detected on the surface of the LAMP-1 fluorophore boundary (Figure 4a).

**Figure 4:**
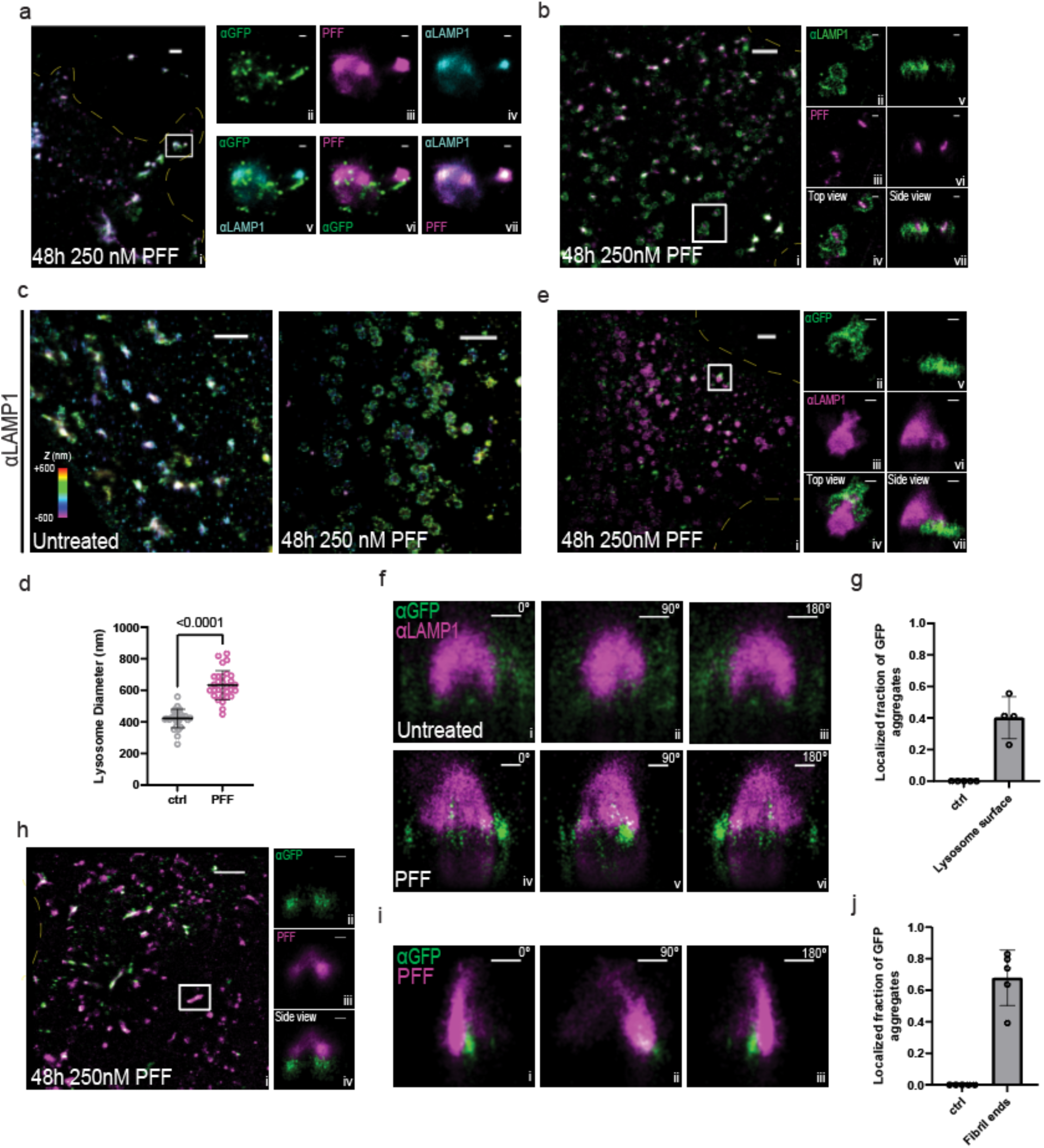
STORM imaging highlights interactions between lysosomes, added tau fibrils, and intracellularly expressed K18.tau-mClover. **a** 3-color STORM image of tau fibrils (magenta), lamp1 (cyan), and tau (green) in a human astrocyte that has been treated with PFF. The full image of the cell is shown on the left, and zoom-ins of the white-boxed region for each channel are shown on the right. The top panels are single-color images of each channel, and the lower panels are two-color merges between channels. **b** Two-color, 3D-STORM images of PFF-treated cells staining PFF (magenta) and lamp1 (green). Side panels show a zoom-in of the white box region. **c** 3D-STORM images of lamp1 in untreated astrocytes (left) and astrocytes incubated with PFF (right). Color encodes axial position (color bar). **d** Histogram of lysosome diameters obtained from the data in **c**. **e** Two-color, 3D-STORM images of PFF-treated cells staining for intracellularly expressed K18.tau-mClover (green) and lamp1 (magenta). Side panels show zoom-in of the white-box region from top (left) and side (right) views. **f** Rotation of lamp1 (magenta) and K18.tau-mClover (green) in untreated (top) and 0N4R fibril-treated cells (bottom). **g** Histogram showing the number of K18.tau-mClover puncta found localized near the surface of the lamp1 lysosome signal. **h** Two-color, 3D-STORM images of 0N4R-treated cells staining intracellularly expressed K18.tau-mClover (green) and 0N4R fibrils (magenta). Side panels show a zoom-in of the white box region from a side view. **i** Rotation of an isolated 0N4R fibril (magenta) and K18.tau-mClover aggregate (green) at 0, 90, and 180°. **j** Histogram of puncta found at the tips of 0N4R fibrils. Scale bars for large images (a,b,c,e,h) = 2 um. Scale bars for smaller panels (a,b,e,f,h,i) = 500 nm.

We next used 3D-STORM to characterize the interactions between K18.tau aggregates with fibrils and lysosomes ^39^. The addition of fibrils to the media increased the diameter of lysosomes by roughly 1.5-fold, from 439 nm to 629 nm (Figure 4b-d). GFP aggregate density was detected primarily on the outside of the LAMP-1 sphere boundary, forming aggregates and extensions projecting away from the center of the lysosome body (Figure 4e-g). GFP aggregates could also be detected on the ends of labeled PFF (Figure 4h-j). Approximately 68% and 40% of the GFP aggregates were localized to fibril ends and lysosome surfaces, respectively (Figure 4g,j).

## Discussion

In the prion-like propagation model for the spread of amyloid in the brain, key steps have been established, including the ability of tau seeds to be endocytosed and to nucleate aggregation of cytosolic tau ^4^. Yet the mechanism for egress from the endosomal pathway to trigger cytosolic tau aggregation has remained mysterious. A nanoscale membrane damage mechanism for tau escape was suggested on the basis of a seeded tau aggregation phenotype of the ESCRT-III subunit CHMP6 ^14^, but the mechanism of action was not resolved. Here, we showed that lysosomes are deacidified following tau uptake, consistent with nanoscale membrane damage. While this could in principle have been explained by V-ATPase inhibition, we further found that the ESCRT-III subunit CHMP4B was recruited to lysosomes under these conditions, diagnostic for nanoscale membrane damage. Moreover, we found that cytosolic tau aggregates at the lysosome membrane surface, and confirmed this event at high spatial resolution using STORM. Collectively, these data are very hard to explain by alternative mechanisms, and establish, at least under the tested conditions in astrocytes, that tau escape occurs via nanoscale lysosomal membrane damage.

The discovery of the role of ESCRTs in lysosomal membrane repair ^22,23^ was followed more recently by characterization of the role of cholesterol and PI(4)P transfer from the ER ^40,41^. Progress in the field has been driven primarily by the use of the model damaging agent LLOME. Recently, defects in the ESCRT pathway in its role as a lysosomal repair machine have been linked to Alzheimer’s disease ^42^. Now we found that tau fibrils are robust inducers of nanoscale lysosomal membrane damage and ESCRT recruitment, which will be important to characterize further as biomedically and physiologically relevant damaging agents.

The nanoscale permeabilization mechanism is radically different from the well-established mechanism for viral pathogen escape from the endolysosomal system, fusion with the endolyosomal membrane ^43^. In contrast to the creation and resolution of a fusion pore by influenza or other viruses, this mechanism implies fibrillar seeds may protrude through the membrane into the cytosol, where they could template nucleation at the membrane surface, which is the predominant mechanism that we see. They could also break off, accounting for a subset of nucleation events that do not appear to be lysosome-adjacent. The observation of lysosomal membrane permeabilization begs the question as to how tau interacts with membranes tightly enough to penetrate them. Unlike synuclein ^44–46^, there have not been extensive reports on how tau interacts with membranes in purified systems ^47^. One possiblity is that, as observed for Huntingtin ^48^, the bulky and rigid fibrils exert mechanical pressure on membranes without necessarily binding to them strongly.

Here we found that lysosomal permeabilization and nanoscale damage, and lysosomal templating by tau, occurs in astrocytes. The data do not establish how common this mechanism is in other systems and relative to other potential mechanisms, such as exosome ^49^ and tunneling nanotube (TNT)-dependent transfer ^50,51^. It will be exciting to explore whether this mechanism is more general, and might apply to the spread of other forms of amyloid in the brain. These data raise questions about the fate of fibrillar cargo taken up by autophagy and lysosomes as a clearance response. If endocytosed fibrils can disrupt lysosomes, presumably fibrils taken up by autophagy could do the same. This highlights the importance of understanding the ESCRT-dependent damage response (Figure 5), and potentially targeting it for upregulation as a therapeutic strategy to prevent spread. This could be of value both from the point of view of restricting cell-to-cell spread, and making autophagic-lysosomal clearance as robust as possible.

**Figure 5:**
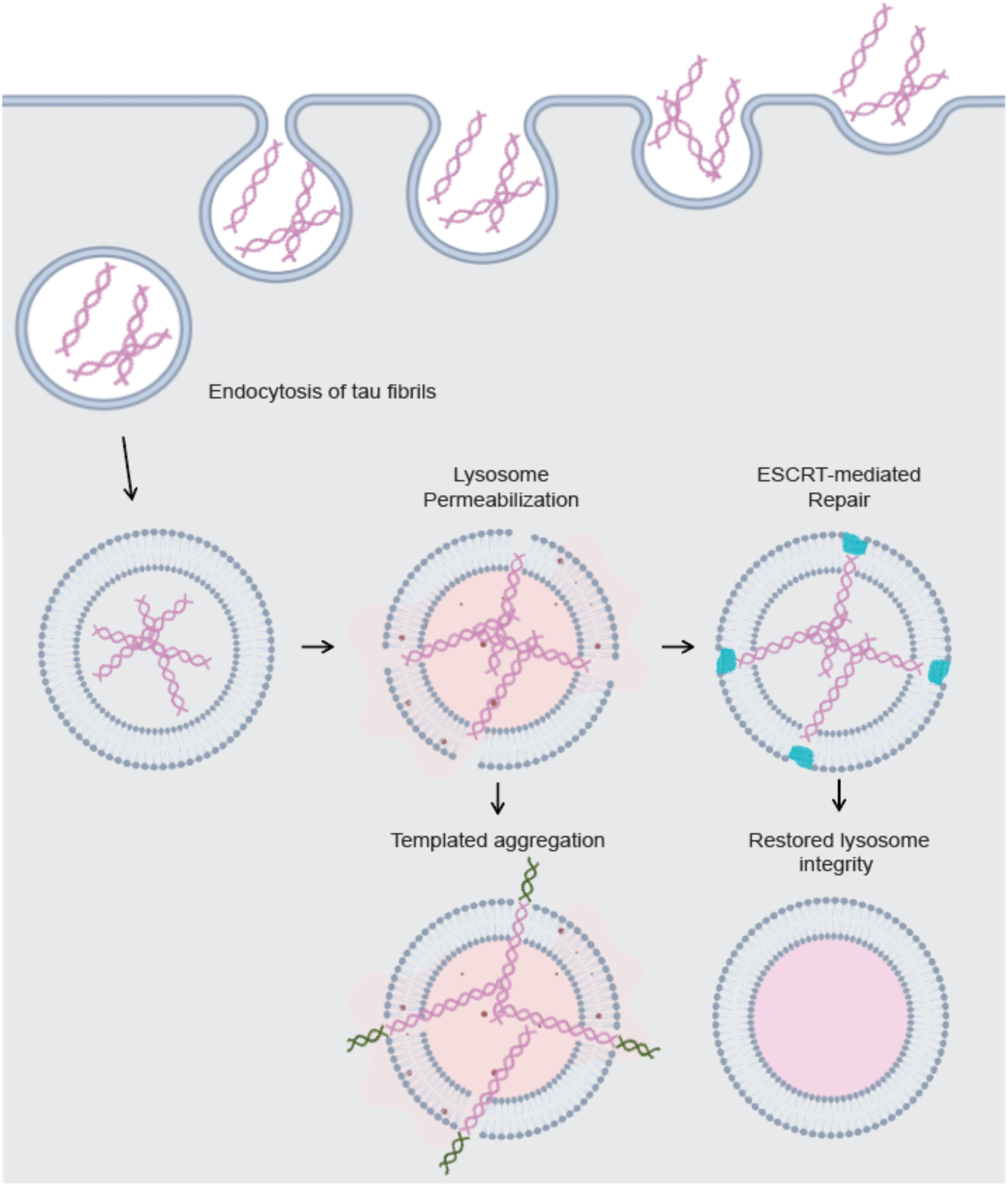
Tau fibrils permeabilize lysosomes but are counteracted by ESCRTs to prevent templated aggregation. Endocytosis of tau fibrils in astrocytes leads to lysosome permeabilization. ESCRTs are recruited to these permeabilized lysosomes to restore membrane integrity and likely promote the continued degradation of intralumenal fibrils. If tau fibril-mediated permeabilization is too extensive to be counteracted by ESCRT proteins, the templated aggregation of cytosolic tau begins on the surface of the lysosome where fibril ends emerge from the membrane.

## Methods

### Cell culture and cell line generation

Human primary astrocytes were obtained from Sciencell (catalog #1800), and cryo-preserved at the UCB Cell Culture Facility by A. Killilea. Human primary astrocytes used for experiments were maintained using Sciencell Astrocyte media (catalog #1801) for no more than 10 passages on plates coated with 2 mg/mL poly-L-Lysine (Sciencell catalog: 0403). HEK293T cells were cultured for no more than 20 passages in DMEM supplemented with 10% fetal bovine serum, Pen/Strep (Life Technologies, catalog number 15140122), and l-glutamine (Life Technologies, catalog number 25030081). All cells were maintained in a copper-lined Heracell VIOS 160i tissue culture incubator (ThermoFischer catalog: 51033574) at 37 °C and 5% CO2 and checked for mycoplasma contamination.

To generate K18.tau-mClover, TMEM192-RFP, pHLARE, CHMP4B-HALO, and Gal3-GFP lentiviruses, 2E6 HEK293T cells were seeded into 10 cm plates and transfected the next day with 45 uL Mirus LT1 transfection reagent (MIR2300) added to a mixture of 5 mg each (15 mg total) of plasmids VSV-G (addgene: 8454), R8.74 (addgene: 22036), and one of the following: K18.tau-mClover (addgene: 133058), TMEM192-RFP (addgene: 134631), pHLARE (addgene: 164478), CHMP4B-HALO (this study), or Gal3-GFP (addgene: 62734) in 1.5 mL Optimem (ThermoFischer catalog: 31985062) according to manufacturer recommendations. Supernatant containing viruses was obtained 3 days post-transfection, clarified by centrifugation at 2000 rpm for 2 minutes, and concentrated 10-fold using Lenti-X concentrator (Takara Bio catalog: 631231). On the day prior to transduction, astrocytes were seeded at a density of 100,000 cells per well into individual wells of a 12 chamber plate (catalog number: 07-200-82). Stable pools of cells expressing desired proteins of interest were obtained by titrating virus concentrate to achieve near 100% expression efficiency, and passaging cells once prior to experiments.

### Production and fibrilization of recombinant 0N4R PFF for cell seeding assays

N-terminal 6x His tagged Human WT 0N4R tau protein is expressed in Rosetta^TM^ 2 (DE3)-competent cells in LB medium supplemented with Ampicillin (100 μg/ml), induced at 0.6 OD with 1 mM isopropyl 1-thio-β-d-galactopyranoside at 37°C for 3 h. After lysis through tip sonication in Buffer A (1x PBS, pH 7.5, 2 mM MgCl_2_, 10 mM EGTA, 1 mM TCEP), supplemented with 0.1 mM PMSF and two EDTA free protease inhibitor tablets (Roche, Basel, Switzerland), the expressed protein was extracted from the supernatant using Pierce^TM^ High-Capacity Ni-IMAC Resin (Thermo Scientific, Waltham, MA) and eluted using lysis buffer supplemented with 300 mM Imidazole pH 7.5. TEV protease was added to the IMAC elute and subjected to overnight dialysis in Buffer B (20 mM MES, pH 6.8, 50 mM NaCl, 1 mM EGTA, 1 mM MgCl_2_, 1 mM TCEP). The dialyzed protein was purified by cation exchange chromatography using 5 ml HiTrap SP (Cytiva, Marlborough, MA) and eluted with a linear gradient of 0 - 1 M NaCl. Fractions containing tau as determined by Coomassie-stained SDS-PAGE were loaded onto the Superdex 200 16/60 column (Cytiva, Marlborough, MA) for gel filtration and eluted using SEC buffer (1x PBS, pH 7.5, 2 mM MgCl_2_, 1 mM TCEP). The concentration of the purified protein was calculated by measuring the absorbance at 280 nm. Finally, the protein was concentrated at approximately 15 μM, snap frozen, and stored at −80°C in 75 μl aliquots.

Aggregation of monomeric tau into PFF was induced by incubating 10 μM tau with 1.3 μM of heparin [Average molecular weight ∼ 16500 g/mol] (Thermo Scientific, Waltham, MA) and shaken at 37 °C overnight at 1200 rpm. The resulting tau fibrils were fluorescently labeled with AlexaFluor 647 NHS-ester dye (Invitrogen, Carlsband, CA) following manufacturer’s protocol. Briefly, NaHCO3 was added to a final concentration of 100 μM to raise the pH of labeling mixture before adding the dye to molar (monomeric) protein to dye ratio of 3:1. This fibril-dye mixture was kept in dark at room temperature for 1 h before removing the unlabeled dye with a Zeba 7k MWCO spin desalting column (Thermo Scientific, Waltham, MA).

For quality control, PFF were negatively stained with 0.8% (w/v) uranyl formate (pH 5.5 - 6.0) on thin amorphous carbon-layered 400-mesh copper grids (Electron Microscopy Services, Hatfield, PA). Five μl of sample was applied to the grid for 1 minute followed by one wash with 6 μl of uranyl formate and a 1 minute incubation with 6 μl of uranyl formate, removed with Whatman^TM^ filter paper (Cytiva, Marlborough, MA). Grids were imaged at room temperature using a Fei Tecnai 12 microscope operating at 120 kV. Images were acquired on a 4000 TVIPS TemCam-F416 camera at ×30000 resulting in a sampling of 3.70 Å/pixel. PFF were typically greater than 10 µm in length but could be broken up into 50 nm fragments using a continual sonication water bath at 37 degrees for 30 minutes, verified by negative stain as above. Sonicated PFF were used in cell seeding experiments by adding the fibrils directly to the media at a final concentration of 250nM (unless otherwise indicated) and monitoring cells for 24-48 hours.

### Monitoring lysosomal pH

pHLARE expressing astrocytes were seeded at a density of 100,000 cells perwell onto poly-L-Lysine (Sigma catalog MILL-A-005-C) coated coverslips in individual wells of a 12 chamber plate. Bafilomycin (Sigma catalog: 19-600-010UG) and PFF treatment was conducted at the indicated concentrations and durations prior to fixing cells with 4% Paraformaldehyde (EMS catalog 15710-S) in DPBS (Corning catalog: 21040CV) for 15 minutes. Fixed coverslips were then mounted onto glass slides using ProLong Gold with DAPI (ThermoFischer catalog: P36935) and left to harden in the dark overnight prior to imaging. Cells were imaged using a using a Nikon A1 confocal microscope with a 63× Plan Apochromat 1.4 numerical aperture objective with identical imaging settings across replicates. Diffuse GFP signal was subtraced and cell images were processed using a median filter of 7 pixels in ImageJ prior to quantification. Total RFP area per cell was then quantified and compared across cells before quantifying the overlapping GFP-positive area using the ImageJ Coloc2 Plugin (https://imagej.net/plugins/coloc-2). Two-way ANOVA was performed in GraphPad Prism 9.0 to compare the effect of Bafilomycin and PFF on lysosomal pH. Coloc2 was additionally used to compare GFP-positive lysosomes to GFP-positive PFF in cells treated with labeled fibrils. Linear regression was performed in GraphPad Prism 9.0.

### Visualization of CHMP4B and Gal3 recruitment to lysosomes

TMEM192-RFP expressing astrocytes were fisrt treated with fluorescently labeled PFF to verify lysosomal accumulation. These cells were additionally transduced with CHMP4B-HALO in parallel experiments for the monitoring of ESCRT responses to PFF treatment. CHMP4B signal was visualized using far-red HALO ligand according to manufacturer recommendations (Promega catalog GA1120). Cells were processed as above for imaging. Number of cells expressing CHMP4B puncta was determined by scanning fields of cells positive for CHMP4B signal and manaully reporting the fraction of cells exhibiting puncta.

TMEM192-RFP expressing astrocytes were also transduced with Gal3-GFP and subjected to the same fibril treatment as CHMP4B-expressing cells. Gal3-GFP recruitment to lysosomes was monitored by TMEM192-RFP colocalization. Gal3-positive TMEM192-RFP area was measured before and after treatment using Coloc2 and normalized to resting levels of Gal3-positive lysosomes. Treated cells were compared to resting cells using an Unpaired t-test in GraphPad Prism 9. Cell images were processed as above for quantification and visualization.

### Live cell imaging and lysosome tracking

Astrocytes expressing K18.tau-mClover and TMEM192-RFP were seeded into poly-L-Lysine coated 35-mm glass bottom dishes (no. P35G-1.5-14-C, MatTek). Cells were treated with unlabeled PFF for 24 and 48 hours prior to conducting live imaging experiments. Timelapse imaging was conducted every 15 seconds for 3 minutes to track lysosomes with adjacent K18.tau aggregates. The centroid of the overlapping fluorescenct region was monitored over time using the ImageJ Particle Tracker Plugin (https://imagej.net/plugins/particle-tracker). The centroid trace was then overlayed on the last frame of the timelapse. Cell images were processed as above for quantification and visualization.

### Super-resolution imaging of PFF-treated astrocytes

For STORM imaging, astrocytes loaded with either unlabeled or AF647 NHS ester-labeled tau fibrils (0N4R PFF, described earlier in methods) were fixed with 3% (vol/vol) paraformaldehyde (Electron Microscopy Sciences, 15714) and 0.1% (vol/vol) glutaraldehyde (Electron Microscopy Sciences, 16365) in DPBS at RT for 20 min, and then washed twice with a freshly prepared 0.1% NaBH_4_ solution followed by three additional washes with DPBS. Cells were then incubated with a solution containing the membrane pore-forming protein Streptoylsin O (SLO) (Sigma catalog: S5265-25KU) sufficient to permeabilize 90% of HeLa cells, as used previously ^52^. SLO-permeabilized cells were washed twice with DPBS followed by incubation with the antibodies of interest, with SLO pores allowing antibodies to pass through cellular membranes while keeping their structure intact. For the two-color images, lamp1 was initially targeted with mouse anti-lamp1 (Santa Cruz Biotech, SC-20011) followed by either donkey anti-mouse CF583R (Biotium catalog: 20794-500UL) for samples loaded with labeled fibrils, or goat anti-mouse AF647 (Thermo-Fischer, A21235) for samples loaded with unlabeled fibrils. K18.tau-mClover was targeted with mouse anti-GFP (Thermo-Fischer, MA5-15256), followed by donkey anti-mouse CF583R (Biotium catalog: 20794-500UL). Three-color STORM used the following labeling scheme: AF647 NHS ester labeled PFF, while donkey anti-rabbit CF680 (Biotium catalog: 20418-50UL) and donkey anti-mouse CF583R (Biotium catalog: 20794-500UL), were used as secondaries for lamp1 (rabbit anti-lamp1, Invitrogen, PA1654A) and GFP (mouse anti-GFP, Thermo-Fischer, MA5-15256), respectively. SLO-permeabilized cells were blocked for 30 minutes with DPBS containing 3% BSA prior to antibody incubation. All primary antibodies were prepared at a dilution of 1:100 and secondary antibodies at a 1:500 dilution, in DPBS containing 3% BSA. Cells were incubated with primary antibodies for 1 hour at room temp, washed 3 times with DPBS between primary and secondary incubations, and incubated with secondary antibodies for 1 hour at room temp in the dark. Residual secondary antibody was removed by three additional washes with DPBS.

Two-color 3D-STORM was performed for samples labeled by Alexa Fluor 647 and CF583R on a homebuilt setup based on a Nikon Eclipse Ti-U inverted microscope, as previously described ^53,54^. The dye-labeled samples were mounted with a Tris⋅HCl–based imaging buffer containing 5% (wt/vol) glucose, 100 mM cysteamine (Sigma-Aldrich, 30070), 0.8 mg/mL glucose oxidase (Sigma-Aldrich, G2133), and 40 µg/mL catalase (Sigma-Aldrich, C30). The sample was sequentially STORM-imaged for Alexa Fluor 647 and CF583R with 647- and 560-nm illuminations at ∼2 kW/cm^2^. The angle of incidence was slightly smaller than the critical angle of total internal reflection, thus illuminating a few micrometers into the sample. A weak (0 to 1 W/cm^2^) 405-nm laser was applied to assist photoswitching. A cylindrical lens was inserted into the optical path for 3D localization ^39^. The resulting stochastic photoswitching of single-molecule fluorescence was recorded using an Andor iXon Ultra 897 EM-CCD camera at 110 frames per second, for ∼80,000 frames per STORM image. The raw STORM data were analyzed using previously described methods ^39^.

Three-color STORM was performed on another homebuilt setup based on a Nikon Eclipse Ti-U inverted microscope, as previously described ^55^. Samples with Alexa Fluor 647 and CF680 were first simultaneously excited under 647-nm illumination and distinguished through ratiometric single-molecule detection in two spectrally split wide-field channels ^56^ using a dichroic mirror (Chroma T685lpxr), and CF583R was subsequently imaged under 560-nm excitation.

## Author contributions

K.R., M.K, and J.H.H. designed research; K.R., T.J, A.M, and S.S. performed research; S.S. contributed new reagents; K.R, T.J, and K.X analyzed data; and K.R. and J.H.H. wrote the paper.

## Acknowledgements

We thank Senthil Kaniyappan for comments on the manuscript. This research was supported by Hoffmann-La Roche as part of the Alliance for Therapies in Neuroscience (J.H.H) and NIH/NIA R01 AG062359 (M.K).

## Competing interest statement

J.H.H. is a co-founder and shareholder of Casma Therapeutics and receives research funding from Genentech and Hoffmann-La Roche. M. K. serves on the Scientific Advisory Boards of Engine Biosciences, Casma Therapeutics, Cajal Neuroscience, Alector, and Montara Therapeutics, and is an advisor to Modulo Bio and Recursion Therapeutics.

